# Lovastatin, not simvastatin, corrects core phenotypes in the fragile X mouse model

**DOI:** 10.1101/430348

**Authors:** Melania Muscas, Susana R. Louros, Emily K. Osterweil

## Abstract

The cholesterol-lowering drug lovastatin corrects neurological phenotypes in animal models of fragile X syndrome (FX), a commonly identified genetic cause of autism and intellectual disability. The therapeutic efficacy of lovastatin is being tested in clinical trials for FX, however the structurally similar drug simvastatin has been proposed as an alternative due to an increased potency and brain penetrance. Here, we perform a side-by-side comparison of the effects of lovastatin and simvastatin treatment on two core phenotypes in the *Fmr1^-/y^* mouse model. We find that while lovastatin normalizes excessive hippocampal protein synthesis and reduces audiogenic seizures (AGS) in the *Fmr1^-/y^* mouse, simvastatin does not correct either phenotype. These results caution against the assumption that simvastatin is a valid alternative to lovastatin for the treatment of FX.

## Introduction

Fragile X syndrome (FX) is a monogenic neurodevelopmental disorder characterized by severe intellectual disability (ID), autism, hypersensitivity to sensory stimulation and epilepsy [1]. FX occurs in 1:4000 males and 1:8000 females, making it one of the most commonly identified genetic causes of autism and ID [1, 2]. The *FMR1* gene mutated in FX encodes Fragile X Mental Retardation Protein (FMRP), which represses mRNA translation in neurons [3, 4]. Studies of the *Fmr1^-/y^* mouse model of FX reveal that excessive cerebral protein synthesis is a major consequence of *Fmr1* deletion [5-9], which can be normalized through antagonism of metabotropic glutamate receptor 5 (mGlu_5_) or the downstream extracellular regulated kinase 1/2 (ERK1/2) MAP kinase signalling pathway [7, 10-13]. These strategies correct multiple neurological phenotypes in the *Fmr1^-/y^* mouse, including an enhanced susceptibility to audiogenic seizures (AGS) [7, 10, 11, 14]. The current challenge is to successfully transition these therapeutic approaches to the clinic.

In previous work we showed that the statin drug lovastatin, currently used for the treatment of high cholesterol in adults and children, reduces ERK1/2 activation and resolves neuropathology in the *Fmr1^-/y^* mouse model [15]. Lovastatin was the first statin drug developed and has been shown to be remarkably effective in lowering cholesterol with minimal side effects [16]. In FX, the therapeutic relevance of lovastatin is in the dampening of ERK1/2 signalling that occurs by reducing the activation of the upstream GTPase Ras [17, 18]. By targeting the same mevalonate pathway that produces cholesterol, lovastatin limits the availability of farnesyl pyrophosphate precursor that is required for the membrane association and activation of Ras [17-19]. By this mechanism, lovastatin has been shown to successfully correct electrophysiological and behavioural phenotypes in the mouse model of Neurofibromatosis Type 1 (NF1), a neurodevelopmental disorder of excess Ras [20].

In the *Fmr1^-/y^* mouse, the reduction of Ras-ERK1/2 by lovastatin ameliorates hippocampal epileptogenesis and neocortical hyperexcitability and significantly reduces the incidence of AGS [15]. Additionally, studies using the *Fmr1^-/y^* rat model show that lovastatin treatment at a juvenile age can prevent the emergence of complex cognitive phenotypes [21]. Based on the positive outcome with lovastatin in *Fmr1^-/y^* animal models, two open-label clinical trials tested the viability of lovastatin for the treatment of FX [22, 23]. Both studies revealed a significant improvement with lovastatin treatment, and a double-blind placebo-controlled trial is ongoing [6].

Interestingly, the availability of lovastatin is not widespread in Europe and is not licensed for use in the UK. Instead, the drug simvastatin has been proposed as an alternative therapeutic. Simvastatin is a structurally similar derivative of lovastatin that is twice as potent, with a daily dose of only 10 mg reducing cholesterol by 25-30% compared to 20 mg of lovastatin [24, 25]. Simvastatin is also more brain penetrant than lovastatin, suggesting it may be a better option for neurological indications [25]. However, simvastatin has not been investigated in the *Fmr1^-/y^* model, and the impact on Ras-ERK1/2 signalling in the brain is not well established. This information is critical, as clinical trials in NF1 have recently shown that lovastatin has a beneficial impact on cognitive function whereas simvastatin does not [26-30].

In this study, we performed a side-by-side comparison of lovastatin and simvastatin to answer the simple but important question of whether there is a similar rescue of pathology in the *Fmr1^-/y^* mouse. We focused on two core phenotypes in the *Fmr1^-/y^* model: excessive protein synthesis and enhanced susceptibility to AGS. Importantly, our results clearly show that lovastatin, but not simvastatin, is effective in reducing ERK1/2 activity and normalizing protein synthesis in the *Fmr1^-/y^* hippocampus. This suggests that simvastatin acts via a different mechanism from lovastatin with respect to ERK1/2-driven protein synthesis in the brain. To examine whether there was a similar impact on pathology, we performed a thorough AGS analysis using multiple doses of simvastatin and two different mouse strains. In all cases, simvastatin failed to reduce AGS in the *Fmr1^-/y^* mouse, whereas lovastatin was significantly effective. This is compelling evidence that simvastatin may not be a suitable replacement for lovastatin with respect to the treatment of FX.

## Materials and methods

### Mice

All mice were naive to drug and behavioral testing prior to experimentation. Mice were group housed with unrestricted food and water access and a 12h light-dark cycle. Room temperature was maintained at 21 ± 2°C. All procedures were performed in accordance with ARRIVE guidelines and regulations set by the University of Edinburgh and the UK Animals Act 1986. *Fmr1^-/y^* mice (Jackson Labs 003025) were maintained on either a C57BL/6J (Charles River) or a mixed C57BL/6J x FVB background (C57BL/6J backcrossed to FVB by 2 generations).

### Metabolic Labeling

Hippocampal slices were prepared from male littermate WT and *Fmr1^-/y^* C57BL/6J mice (P25-32), in an interleaved fashion, with the experimenter blind to genotype as described previously [11]. Briefly, mice were anaesthetized with isoflurane and the hippocampus was rapidly dissected in ice-cold ACSF (124 mM NaCl, 3 mM KCl, 1.25 mM NaH_2_PO_4_, 26 mM NaHCO_3_, 10 mM dextrose, 1 mM MgCl_2_ and 2 mM CaCl_2_, saturated with 95% O_2_ and 5% CO_2_). Slices (500 µm thick) were prepared using a Stoelting Tissue Slicer and transferred into 32.5°C ACSF (saturated with 95% O_2_ and 5% CO_2_) within 5 min. Slices were incubated in 32.5°C ACSF for 4 hr to allow for recovery of protein synthesis then transferred to ACSF containing 25 μM Actinomycin D (Tocris) plus either vehicle (0.05% DMSO in ddH_2_O), 50 μM lovastatin active form (CAS 75225-50-2; Calbiochem Merck Millipore), or 0.1-5 μM simvastatin active form (CAS 101314-97-0; Cayman Chemical) for 30 minutes. To measure new protein synthesis, slices were then transferred to fresh ACSF with 10 µCi/ml ^35^S-Met/Cys (Perkin Elmer) containing vehicle or drug for another 30 min.

After labeling, slices were homogenized in ice-cold buffer (10 mM HEPES pH 7.4, 2 mM EDTA, 2 mM EGTA, 1% Triton X-100, protease inhibitors and phosphatase inhibitors) and incubated in trichloroacetic acid (TCA: 10% final) for 10 min on ice before being centrifuged at 16,000 rpm for 10 min. The pellet was washed in ice-cold ddH_2_O and re-suspended in 1 N NaOH until dissolved, and the pH was re-adjusted to neutral using 0.33 N HCl. Triplicates of each sample were subjected to scintillation counting and protein concentration assay kit (BioRad). Counts per minute (CPM) were divided by protein concentration, and this was normalized to the CPM from the ACSF used for incubation. For display purposes, example slice homogenates were resolved on SDS-PAGE gels, transferred to nitrocellulose and exposed to a phosphorimaging screen (GE Healthcare). Phosphorimages were acquired using a Typhoon scanner (GE Healthcare) and compared to total protein staining of the same membrane.

### Immunoblotting

Samples were resolved on SDS-PAGE gels, transferred to nitrocellulose and stained for total protein with the Memcode Reversible staining kit (Pierce). Membranes were later blocked with 5% BSA in TBS + 0.1% Tween-20 and incubated in primary antibody overnight at 4°C (Cell Signaling Technology: phospho-ERK1/2 (Thr202/Tyr204), 1:2000 (#9106), ERK1/2 1:2000 (#9102), phospho-p70S6K (Thr389) 1:1000 (#9234), p70S6K 1:1000 (#2708). Membranes were then washed, incubated with HRP-conjugated secondary antibodies for 30 min (Cell Signaling), and developed with Clarity ECL (BioRad). To compare phopho- to total for each target in the same lane, membranes developed for phospho (i.e., p-ERK1/2) were stripped and re-probed for total (i.e., ERK1/2). Densitometry was performed on scanned blot films using ImageStudio Lite software. Phosphorylation of target proteins was calculated as a ratio of phospho- to total. To correct for blot-to-blot variance, each signal was normalized to the average signal of all lanes on the same blot. All gels were loaded and analyzed by experimenter blind to genotype and treatment.

### Audiogenic Seizures

Experiments were performed as previously described [15, 31]. Test cohorts were counterbalanced for genotype and treatment. Naive WT and *Fmr1^-/y^* male P18-29 mice bred on a C57BL/6J or mixed C57BL/6J x FVB background were weighed and injected intraperitoneally (i.p.) with 3 mg/kg simvastatin prodrug (CAS 79902-63-9), 50 mg/kg simvastatin active form (CAS 101314-97-0), or 100 mg/kg lovastatin active form (CAS 75225-50-2) or respective vehicle (3, 20, or 50 % DMSO + 10% Tween-80 in PBS). Animals were then transferred to a quiet (< 60 dB ambient sound) room for 1 hr. For testing, animals were moved to a transparent test chamber equipped with speakers and a webcam and allowed to habituate for 1 min. Audiogenic stimulation (recorded sampling of a modified personal alarm) was passed through an amplifier and 2 X 50-Watt speakers (KRK Rokit RP5 G3 Active Studio Monitor) to produce a stimulus of > 130 dB for 2 min. A decibel meter was placed at a standard distance from the speakers to ensure a stable emission of sound throughout each session. Incidence and severity of seizures was scored and video files for each session were saved. Stages of AGS severity were assigned according to previous work as follows: (1) wild running (WR; pronounced, undirected running and thrashing), (2) clonic seizure (violent spasms accompanied by loss of balance), or (3) tonic seizure (loss of movement and postural rigidity in limbs and tail). Any animal that reached tonic seizure was immediately humanely euthanized. All injections, testing and scoring was performed with the experimenter blind to genotype and treatment.

### Statistics

Statistical testing was performed using GraphPad Prism software. For biochemistry experiments, outliers > 2 SD from the mean were removed and significance determined by repeated measures two-way ANOVA and post hoc Sidak multiple comparisons test. For AGS experiments, significance was determined by Fisher’s exact test. Results of all statistical analyses are reported in detail in the figure legends.

## Results

### Lovastatin, but not simvastatin, normalizes excessive protein synthesis in the *Fmr1^-/y^* hippocampus

In previous work we showed that lovastatin normalizes excessive protein synthesis in the *Fmr1^-/y^* hippocampus through reduction of Ras-ERK1/2 activation, which corrects epileptogenic phenotypes [15]. To examine whether the same effect is seen with simvastatin, we utilized a metabolic labeling assay in hippocampal slices designed to assess protein synthesis in an intact preparation under physiological conditions [11]. Hippocampal slices were prepared from juvenile WT and *Fmr1^-/y^* littermates, blind to genotype, and allowed to recover in oxygenating ACSF. Following this, slices were pre-incubated with Actinomycin D to block transcription, and new protein synthesis was labelled through incorporation of ^35^S-labeled methionine/cysteine mix (**Figure 1A**).

**Figure 1.**
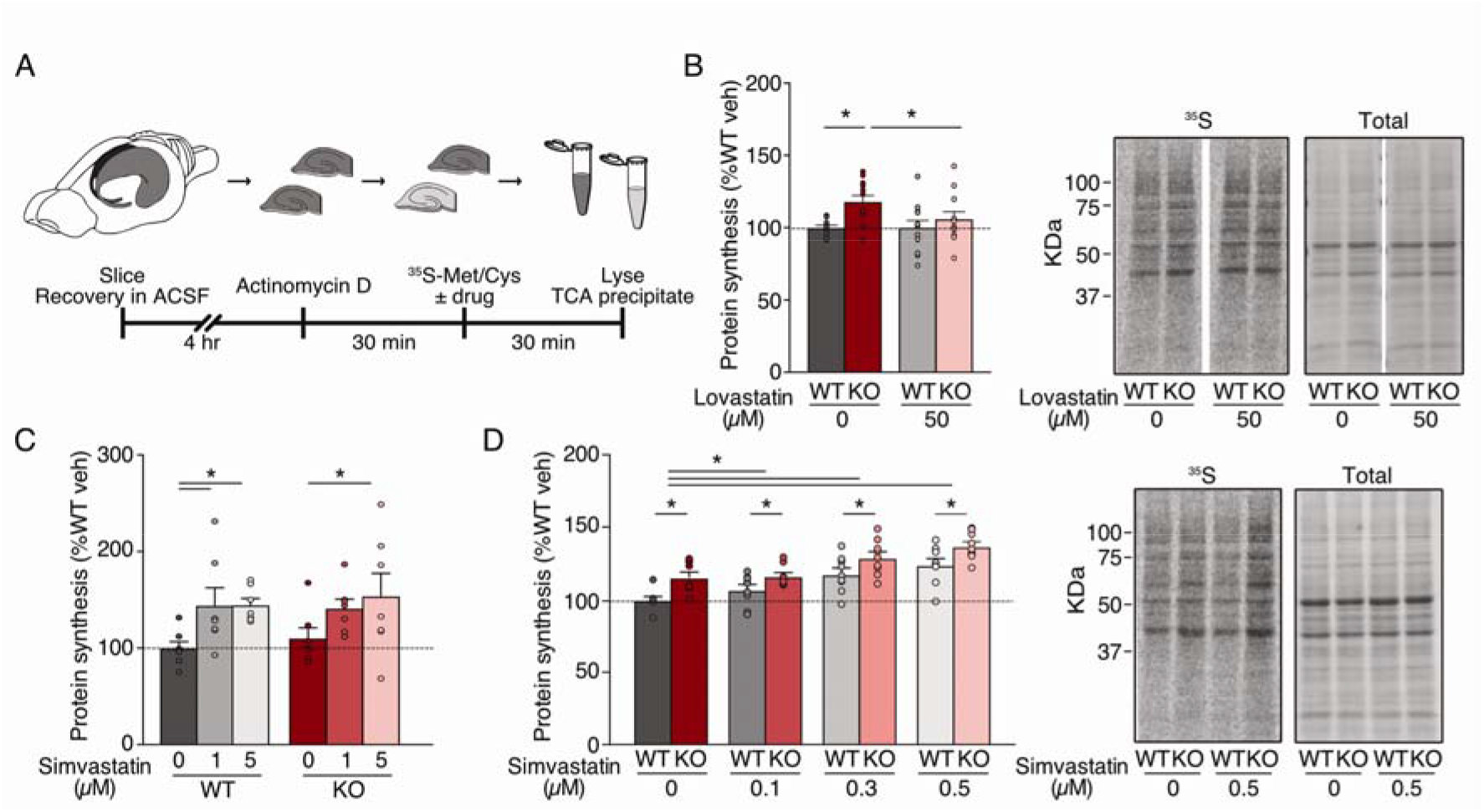
Simvastatin exaggerates excessive protein synthesis in the *Fmr1^-/y^* hippocampus. Slices were prepared from WT and *Fmr1^-/y^* hippocampi and incubated in vehicle, lovastatin or simvastatin at different concentrations. **(A)** Schematic shows time course for metabolic labelling experiments of hippocampal slices. **(B)** Lovastatin significantly decreases protein synthesis in *Fmr1^-/y^* slices to WT levels (WT vehicle = 100 ± 1.48%, WT lova = 100.06 ± 4.87%, KO vehicle = 117.97 ± 4.27%, KO lova = 106.04 ± 4.93%; ANOVA genotype *p = 0.0106; Sidak WT veh vs KO veh *p = 0.0032, WT lova vs KO lova p = 0.3516, KO veh vs KO lova *p = 0.0368; n = 12). **(C)** Simvastatin raises protein synthesis in both WT and *Fmr1^-/y^* slices at 1-5 μM (WT vehicle = 100 ± 2.74%, WT 1 μM = 144.15 ± 14.96%, WT 5 μM = 146.47 ± 6.98%, KO vehicle = 110.60 ± 7.48%, KO 1 μM = 144.56 ± 13.05%, KO 5 μM = 154.90 ± 21.31%; ANOVA treatment *p = 0.0125; Sidak WT veh vs 1μM *p = 0.0371, WT veh vs 5 μM *p = 0.0279, KO veh versus 5 μM *p = 0.0364; n = 7). **(D)** Simvastatin raises protein synthesis at 0.1-0.5 μM, exaggerating the excessive protein synthesis phenotype (WT vehicle = 100 ± 2.21%, WT 0.1 μM = 106.99 ± 3.51%, WT 0.3 μM = 117.79 ± 4.08%, WT 0.5 μM = 124.13 ± 4.23%, KO vehicle = 115.61 ± 3.48%, KO 0.1 μM = 116.52 ± 2.21%, KO 0.3 μM = 129.15 ± 3.99%, KO 0.5 μM = 137.01 ± 3.08%; ANOVA treatment *p < 0.0001, genotype *p = 0.0068; Sidak WT veh vs 0.3 μM *p = 0.0002, WT veh vs 0.5 μM *p < 0.0001, KO veh vs 0.3 μM *p = 0.0035, KO veh vs 0.5 μM *p < 0.0001, WT veh vs KO veh *p = 0.0005, WT 0.1μM vs KO 0.1μM *p = 0.0406, WT 0.3 μM vs KO 0.3μM *p = 0.0115, WT 0.5 μM vs KO 0.5 μM *p = 0.0038; n = 9). Representative samples were run on SDS PAGE gels and transferred to membranes. Example phosphorimages of ^35^S-labelled proteins and total protein staining of the same membrane are shown. Error bars = SEM. N = littermate pairs.

Previous experiments tested a range of 10-50 µM lovastatin and showed that 50 µM was effective in normalizing protein synthesis in the *Fmr1^-/y^* hippocampus [15]. To ensure that we could recapitulate these results, we measured protein synthesis in WT and *Fmr1^-/y^* slices ± 50 µM lovastatin ( **Figure 1B**). As expected, our experiments revealed a significant correction of excessive protein synthesis with lovastatin in the *Fmr1^-/y^* mouse (WT veh = 100 ± 1.48%, WT lova = 100.06 ± 4.87%, KO veh = 117.97 ± 4.27%, KO lova = 106.04 ± 4.93%; WT vs KO veh p = 0.0032, KO veh vs lova p = 0.0368; n = 12). Next, we tested the efficacy of simvastatin using the same assay system. Based on the increased potency of simvastatin, and previous studies of simvastatin in neurons, we tested a range of 1-5 µM [32-34]. Interestingly, we find that simvastatin treatment not only fails to reduce protein synthesis in the *Fmr1^-/y^* hippocampus, it causes a significant increase in both WT and *Fmr1^-/y^* slices at 1-5 µM (WT vehicle = 100 ± 2.74%, WT 1 μM = 144.15 ± 14.96%, WT 5 μM = 146.47 ± 6.98%, KO vehicle = 110.60 ± 7.48%, KO 1 μM = 144.56 ± 13.05%, KO 5 μM = 154.90 ± 21.31%; WT veh vs 1μM p = 0.0371, WT veh vs 5 μM p = 0.0279, KO veh vs 5 μM p = 0.0364; n = 7) (**Figure 1C)**.

This puzzling increase in protein synthesis led us to wonder whether a reduced concentration of simvastatin might be more appropriate. To test this, we exposed slices to vehicle or simvastatin at concentrations of 0.1-0.5 µM. Surprisingly, we find that even at these lower concentrations simvastatin causes a dose-dependent increase in protein synthesis, worsening the *Fmr1^-/y^* phenotype (WT veh = 100 ± 2.21%, WT 0.1 μM = 106.99 ± 3.51%, WT 0.3 μM = 117.79 ± 4.08%, WT 0.5 μM = 124.13 ± 4.23%, KO veh = 115.61 ± 3.48%, KO 0.1 μM = 116.52 ± 2.21%, KO 0.3 μM = 129.15 ± 3.99%, KO 0.5 μM = 137.01 ± 3.08%; WT veh vs 0.3 μM p = 0.0002, WT veh vs 0.5 μM p < 0.0001, KO veh vs 0.3 μM p = 0.0035, KO veh vs 0.5 μM p < 0.0001; n = 9) (**Figure 1D**). These results show that unlike lovastatin, simvastatin does not correct excessive protein synthesis in the *Fmr1^-/y^* hippocampus.

### Lovastatin, but not simvastatin, reduces ERK1/2 activation

Given the differential efficacy of lovastatin and simvastatin on excessive protein synthesis in the *Fmr1^-/y^*, we wondered whether these compounds acted differently on translation control signaling pathways. Two major intracellular signaling pathways implicated in synaptic protein synthesis are the ERK1/2 pathway that stimulated translation initiation through eukaryotic initiation factor 4E (eIF4E), and the mammalian target of rapamycin complex 1 (mTORC1) signaling pathway that activates p70 S6 kinase (p70S6K) to phosphorylate ribosomal protein S6 [35] (**Figure 2A**). These two pathways lie downstream of the GTPases Ras and Rheb, both of which are regulated by farnesylation [36]. The reduced activation of Ras-ERK1/2 signaling upon lovastatin treatment has been documented in several contexts, including hippocampal slices [15, 17, 19, 20]. To confirm this, we incubated slices in vehicle or 50 µM lovastatin and performed quantitative immunoblotting for phosphorylated (p-) ERK1/2 (**Figure 2B**). Our results confirm that 50 µM lovastatin significantly reduces p-ERK1/2 in *Fmr1^-/y^* slices as previously reported (WT veh = 100 ± 4.32%, WT lova = 99.28 ± 4.42%, KO veh = 91.83 ± 4.74%, KO lova = 76.28 ± 3.76%; KO veh vs lova p = 0.0048; n = 19). To examine whether simvastatin had a similar impact on ERK1/2 signaling, we performed the same immunoblotting analysis on slices exposed to vehicle or 0.1-0.5 µM simvastatin. In contrast to lovastatin, our results show that simvastatin has no significant impact on p-ERK1/2 in either WT or *Fmr1^-/y^* slices at any dose tested (WT veh = 100 ± 4.51%, WT 0.1 μM = 102.87 ± 3.42%, WT 0.3 μM = 108.45 ± 4.10%, WT 0.5 μM = 101.01% ± 2.09%, KO veh = 105.63 ± 4.97%, KO 0.1 μM = 98.94 ± 4.46%, KO 0.3 μM = 94.71 ± 4.53%, KO 0.5 μM = 106.93 ± 3.65%; n = 11) (**Figure 2C**).

**Figure 2.**
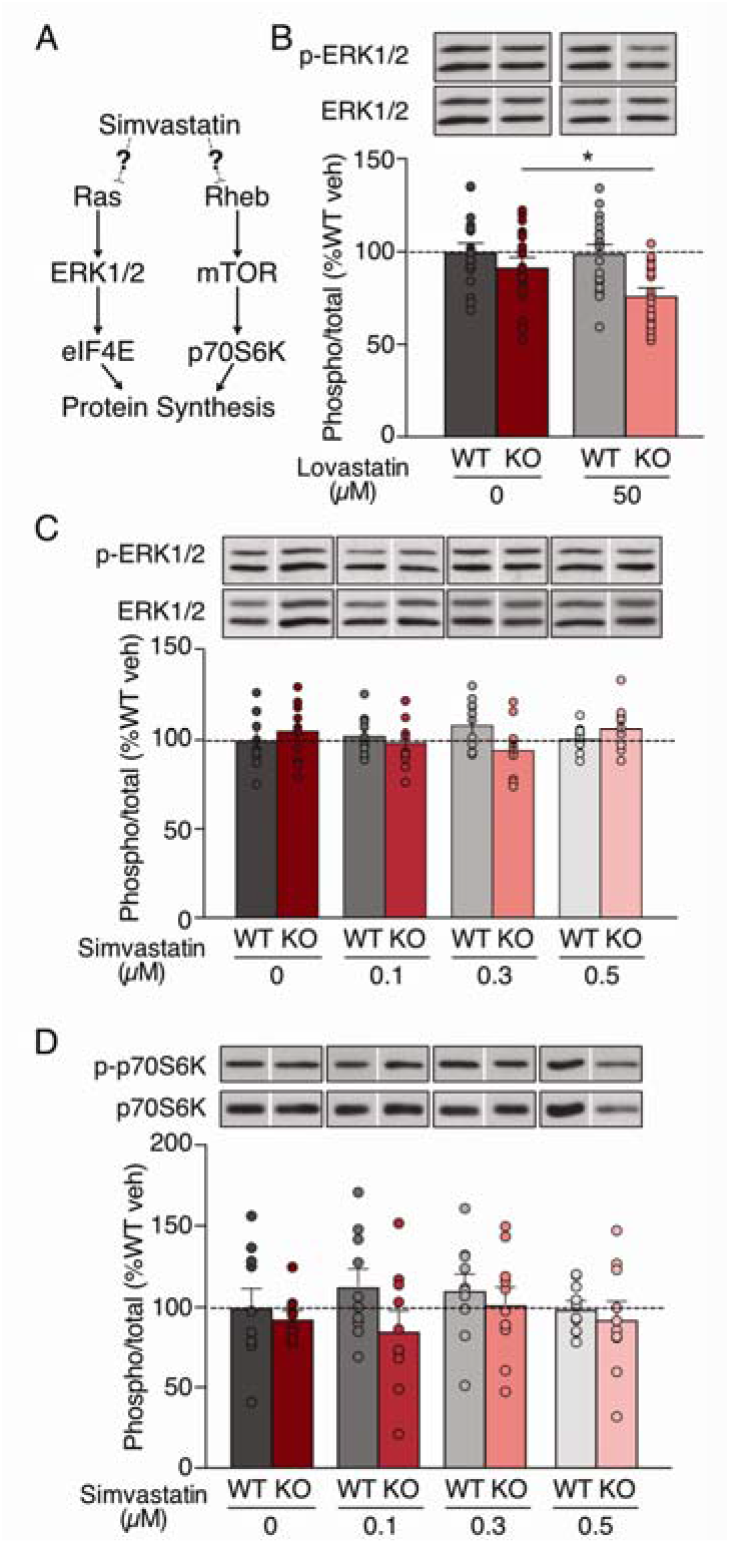
Simvastatin does not reduce ERK1/2 or mTORC1 activation in the *Fmr1^-/y^* hippocampus. **(A)** Diagram shows the potential impact of simvastatin on Ras-ERK1/2 and Rheb-mTOR signaling pathways. **(B)** *Fmr1^-/y^* slices incubated with 50 µM lovastatin show a significant reduction in ERK1/2 phosphorylation (WT vehicle = 100 ± 4.32%, WT lova = 99.28 ± 4.42%, KO vehicle = 91.83 ± 4.74%, KO lova = 76.28 ± 3.76%; ANOVA genotype *p = 0.0146; Sidak KO veh vs KO lova *p = 0.0048; n = 19). **(C)** Simvastatin treatment does not reduce ERK1/2 phosphorylation in *Fmr1^-/y^* or WT slices (WT vehicle = 100 ± 4.51%, WT 0.1 μM = 102.87 ± 3.42%, WT 0.3 μM = 108.45 ± 4.10%, WT 0.5 μM = 101.01 ± 2.09%, KO vehicle = 105.63 ± 4.97%, KO 0.1 μM = 98.94 ± 4.46%, KO 0.3 μM = 94.71 ± 4.53%, KO 0.5 μM = 106.93 ± 3.65%; ANOVA treatment p = 0.8761, genotype p = 0.7010; n = 11). **(D)** Simvastatin treatment does not reduce phosphorylation of p70S6K in WT or *Fmr1^-/y^* slices (WT vehicle = 100 ± 11.14%, WT 0.1 μM = 112.94 ± 10.25%, WT 0.3 μM = 110.66 ± 9.47%, WT 0.5 μM = 98.89 ± 4.72%, KO vehicle = 92.87 ± 4.49%, KO 0.1 μM = 85.37 ± 11.82%, KO 0.3 μM = 101.71 ± 10.37%, KO 0.5 μM = 92.53 ± 10.64%; ANOVA treatment p = 0.6206, genotype p = 0.2860, n = 10). Representative bands were cropped from original blots as indicated by blank spaces. Error bars = SEM. N = littermate pairs.

In addition to Ras, the farnesylation inhibition resulting from reduction of mevalonate upstream cholesterol has been reported to reduce the activation of Rheb [37, 38]. Although our previous study with lovastatin showed no effect of lovastatin on mTORC1 activation as assessed by phosphorylation of p70S6K, we wondered whether simvastatin had an observable impact on this pathway. To investigate, we immunoblotted for p-p70S6K in WT and *Fmr1^-/y^* slices treated with 0.1-0.5 µM simvastatin. Our results show that p70S6K activation is unchanged in slices treated with 0.1-0.5 µM simvastatin (WT veh = 100 ± 11.14%, WT 0.1 μM = 112.94 ± 10.25%, WT 0.3 μM = 110.66 ± 9.47%, WT 0.5 μM = 98.89 ± 4.72%, KO veh = 92.87 ± 4.49%, KO 0.1 μM = 85.37% ± 11.82%, KO 0.3 μM = 101.71% ± 10.37%, KO 0.5 μM = 92.53% ± 10.64%; n = 10) (**Figure 2D**). These experiments show that unlike lovastatin, simvastatin does not suppress the activation of ERK1/2, and it has no effect on mTORC1 activation.

### Lovastatin, but not simvastatin, corrects the AGS phenotype in the *Fmr1^-/y^* mouse

Our work *in vitro* shows that simvastatin does not correct the ERK1/2-stimulated excess in protein synthesis in the *Fmr1^-/y^* hippocampus, suggesting that it may not have the same efficacy as lovastatin in ameliorating pathological phenotypes. To directly test this, we performed a side-by-side analysis of the effect of lovastatin versus simvastatin on the incidence of AGS in the *Fmr1^-/y^* mouse. The AGS phenotype is one of the most robust behavioral phenotypes seen in the *Fmr1^-/y^* mouse, and it models the epilepsy observed in FX patients [39, 40]. Several previous studies have used AGS as a benchmark for determining the efficacy of potential treatment strategies, consistently finding a positive correlation between treatment efficacy at reducing seizure incidence and correction of other pathologies [7, 11, 15, 41-43].

In previous work, 1 mg/kg simvastatin was shown to sufficiently reduce epileptogenic activity and neurotoxicity in a kainic acid (KA) rat model of epilepsy [44]. This suggested simvastatin might be effective in reducing AGS in the *Fmr1^-/y^* mouse even when protein synthesis is not normalized. To investigate, we tested the effect of 3 mg/kg simvastatin on AGS incidence and severity in *Fmr1^-/y^* and littermate WT mice bred on a C57BL/6J background as described in Materials and Methods. Animals were injected with vehicle or simvastatin with the experimenter blind to genotype and treatment, and then left in a quiet environment for 1 hour. To induce AGS, animals were transferred to a test chamber and exposed to a 2-minute digitized sampling of a personal alarm passed through 50-Watt speakers a level of greater than 130 decibels. Seizures were recorded at increasing levels of severity as: 1 - wild running (uncontrolled and undirected running), 2 - clonic seizure (loss of balance with violent spasms on all limbs), and 3 - tonic seizure (loss of balance with postural rigidity in limbs and tail) (**Figure 3A**).

**Figure 3.**
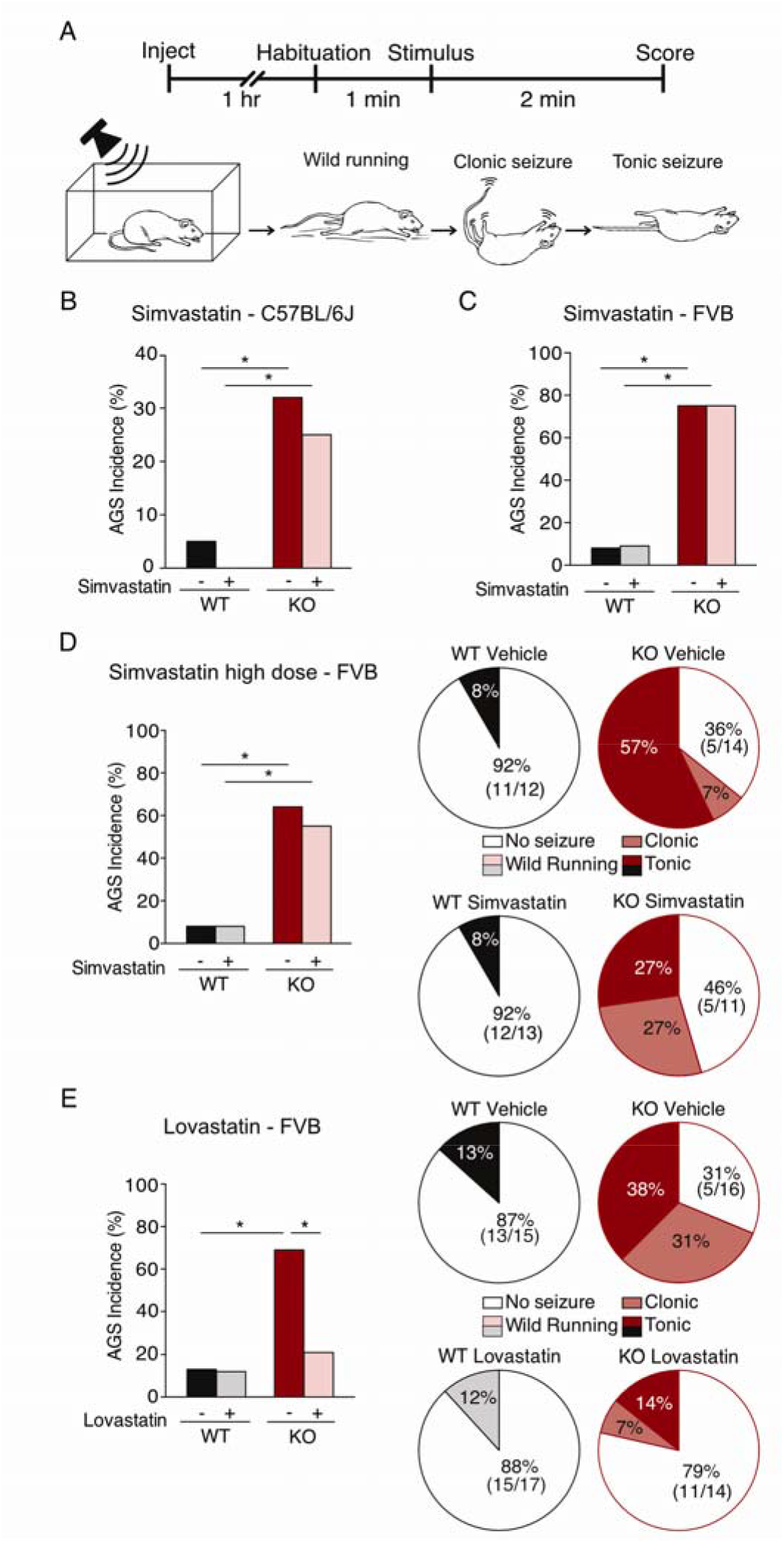
Simvastatin does not correct AGS in the *Fmr1^-/y^* mouse. *Fmr1^-/y^* and littermate WT mice were injected i.p. with vehicle, simvastatin or lovastatin and tested for AGS. **(A)** Schematic shows the experimental timeline and scoring system for AGS testing. **(B)** Injection of 3 mg/kg simvastatin does not reduce AGS incidence in *Fmr1^-/y^* mice bred on a C57BL/6J background (WT veh 4%, WT simva 0%, KO veh 30%, KO simva 22%; Fisher’s exact test WT vs KO veh *p = 0.0279, WT vs KO simva *p = 0.0250, KO veh vs simva p = 0.7570). AGS severity is also not reduced (KO veh: wild running 5/27, clonic 3/27, tonic 0/27; KO simva: wild running 2/27, clonic 4/27, tonic 0/27). **(C)** 3 mg/kg simvastatin does not reduce incidence of AGS in *Fmr1^-/y^* mice bred on a C57BL/6J x FVB background (WT veh 8%, WT simva 9%, KO veh 75%, KO simva 75%; Fisher’s exact test WT vs KO veh *p = 0.0028, WT vs KO simva *p = 0.0028, KO veh vs simva p = 1.000). AGS severity is also not reduced (KO veh: wild running 1/12, clonic 5/12, tonic 3/12; KO simva: wild running 3/12, clonic 2/12, tonic 4/12). **(D**) 50 mg/kg active simvastatin does not reduce AGS incidence in *Fmr1^-/y^* C57BL/6J x FVB mice (WT veh 8%, WT simva 8%, KO veh 64%, KO simva 55%; Fisher’s exact test WT vs KO veh *p = 0.0053, WT vs KO simva *p = 0.0233, KO veh vs simva p = 0.6968). 50 mg/kg simvastatin does not reduce AGS severity in *Fmr1^-/y^* mice (KO veh: wild running 0/14, clonic 1/14, tonic 8/14; KO simva: wild running 0/11, clonic 3/11, tonic 3/11). **(E)** Injection of lovastatin (100 mg/kg active form) significantly reduces the incidence of AGS in *Fmr1^-/y^* C57BL/6J x FVB mice (WT veh 13%, WT lova 12%, KO veh 69%, KO lova 21%; Fisher’s exact test WT vs KO veh *p = 0.0032, KO veh vs lova *p = 0.0136, WT veh vs KO lova p = 0.6513). Lovastatin reduces severity of AGS in *Fmr1^-/y^* mice versus vehicle (KO veh: wild running 0/16, clonic 5/16, tonic 6/16; KO lova: wild running 0/14, clonic 1/14, tonic 2/14).

Our results show that vehicle treated *Fmr1^-/y^* mice exhibit a higher incidence of AGS versus WT littermates (WT veh 4%, KO veh 30%; WT vs KO veh p = 0.0279). However, in contrast to our previous work with lovastatin, we found that simvastatin injection had no significant effect on the incidence of AGS in *Fmr1^-/y^* mice (WT simva 0%, KO simva 22%; WT vs KO simva p = 0.0250, KO veh vs simva p = 0.7570), nor did it reduce AGS severity (KO veh: wild running 5/27, clonic 3/27, tonic 0/27; KO simva: wild running 2/27, clonic 4/27, tonic 0/27) (**Figure 3B**). Although the AGS phenotype is seen in *Fmr1^-/y^* mice bred on multiple mouse background strains, the FVB background strain exhibits a higher incidence of AGS that persists into adulthood [41, 45]. To ensure that the failure to observe a reduction of seizures with simvastatin was not due to a mouse strain-dependent effect, we performed additional AGS experiments using *Fmr1^-/y^* C57BL/6J mice backcrossed two generations to FVB. Consistent with other studies, we found that vehicle treated *Fmr1^-/y^* mice showed a higher incidence of AGS with introduction of the FVB strain (**Figure 3C**) [41, 45]. However, 3 mg/kg simvastatin remained ineffective in reducing the AGS phenotype in *Fmr1^-/y^* mice even on the C57BL/6J x FVB background (WT veh 8%, WT simva 9%, KO veh 75%, KO simva 75%; WT vs KO veh p = 0.0028, WT vs KO simva p = 0.0028, KO veh vs simva p = 1.000). AGS severity was also not reduced by simvastatin (KO veh: wild running 1/12, clonic 5/12, tonic 3/12; KO simva: wild running 3/12, clonic 2/12, tonic 4/12) (**Figure 3C**). Hence, 3 mg/kg simvastatin does not correct the AGS phenotype in *Fmr1^-/y^* mice on either background strain.

A broad range of simvastatin doses has been used to examine the potential impact on KA-induced seizures and other neurological phenotypes in mice, with the 50 mg/kg being the highest dose tested [46, 47]. An equipotent 100 mg/kg dose of lovastatin was previously shown to correct AGS in adult *Fmr1^-/y^* FVB mice, and we wondered whether the higher 50 mg/kg dose of simvastatin would be similarly effective [15]. Additional AGS experiments were performed on *Fmr1^-/y^* and WT C57BL/6J x FVB mice injected with vehicle or 50 mg/kg simvastatin. To ensure a maximal potency, we injected the active form of simvastatin that does not require hydrolyzation from a prodrug form [25]. Our results show that even at this higher dose, simvastatin does not reduce AGS incidence in *Fmr1^-/y^* mice (WT veh 8%, WT simva 8%, KO veh 64%, KO simva 55%; WT vs KO veh p = 0.0053, WT vs KO simva p = 0.0233, KO veh vs simva p = 0.6968), nor does it reduce AGS severity (KO veh: wild running 0/14, clonic 1/14, tonic 8/14; KO simva: wild running 0/11, clonic 3/11, tonic 3/11) (**Figure 3D**). In contrast, *Fmr1^-/y^* C57BL/6J x FVB mice injected with 100 mg/kg lovastatin showed a significant reduction in AGS versus vehicle-treated mice (WT veh 13%, WT lova 12%, KO veh 69%, KO lova 21%; WT vs KO veh p = 0.0032, KO veh vs lova p = 0.0136, WT veh vs KO lova p = 0.6513) (**Figure 3E**). Together, these results show that lovastatin, not simvastatin, is effective in ameliorating AGS in the *Fmr1^-/y^* model.

## Discussion

The promising results using lovastatin in FX have led to the suggestion that simvastatin may be similarly effective. In this study, we performed a thorough analysis of two core phenotypes in the *Fmr1^-/y^* mouse model to test the prediction that simvastatin can be used in place of lovastatin. Our results show that simvastatin not only fails to correct excessive protein synthesis in the *Fmr1^-/y^* hippocampus, it worsens this phenotype (**Figure 1**). Moreover, simvastatin does not reduce ERK1/2 activation in the *Fmr1^-/y^* hippocampus, suggesting it does not share the same mechanism as lovastatin in reducing Ras activity (**Figure 2**). Importantly, analysis of the AGS phenotype in the *Fmr1^-/y^* mouse shows that simvastatin does not reduce seizures in either of the two mouse strains tested, even at high doses (**Figure 3**). Together, these results suggest that simvastatin is not an effective replacement for lovastatin with respect to the treatment of FX.

Although we propose the beneficial effect of lovastatin stems from the inhibition of ERK1/2-driven protein synthesis, it is important to note that statins are capable of affecting several biochemical pathways. Beyond the canonical impact on cholesterol biosynthesis, statins also decrease isoprenoid intermediates including farnesyl and geranylgeranyl pyrophosphates that regulate membrane association for many proteins including the small GTPases Ras, Rho and Rac [18, 46, 48, 49]. The increase in protein synthesis seen with simvastatin could be linked to altered posttranslational modification of these or other proteins. Indeed, although we see no change in mTORC1-p70S6K signaling, other studies have shown an activation of the PI3 kinase pathway that could be contributing to this effect [32]. However, our comparison of lovastatin and simvastatin shows that there is a clear difference in the correction of pathology in the *Fmr1^-/y^* model, suggesting that the impact on ERK1/2 is an important factor in terms of pharmacological treatment for FX.

There are many reasons why statins would be an attractive option for treating neurodevelopmental disorders such as FX. They are widely prescribed worldwide for the treatment of hypercholesterolemia and coronary heart disease [50], and safely used for long-term treatment in children and adults [46]. However, our study suggests that care should be taken when considering which statin should be trialed for the treatment of FX and other disorders of excess Ras. Although the effect of different statins on cholesterol synthesis has been well documented, the differential impact on Ras-ERK1/2 signaling is not well established. We show here that, contrary to lovastatin, simvastatin fails to inhibit the Ras-ERK1/2 pathway in the *Fmr1^-/y^* hippocampus, exacerbates the already elevated protein synthesis phenotype, and does not correct the AGS phenotype. These results are significant for considering future clinical trials with lovastatin or simvastatin for FX or other disorders of excess Ras. Indeed, clinical trials using simvastatin for the treatment of NF1 have shown little promise, while trials with lovastatin show an improvement in cognitive deficits [28-30]. We suggest that simvastatin could be similarly ineffective in FX and may not be a suitable substitute for lovastatin in further clinical trials.

## Acknowledgements

We would like to acknowledge the contributions of all members of the Osterweil lab, with special thanks to Sophie Thomson, Sang Seo, Steph Barnes, and Caoimhe Kirby. We also thank Peter Kind and Mike Cousin for helpful insights and advice.

## Author Contributions

MM and SRL performed all biochemistry and audiogenic seizure experiments. EO, MM and SRL conceptualized the study and prepared the manuscript.

## References

1. Lozano R, Rosero CA, and Hagerman RJ. Fragile X spectrum disorders. Intractable Rare Dis Res. 2014; 3(4): p. 134–46.

2. Hagerman RJ, Berry-Kravis E, Kaufmann WE, Ono MY, Tartaglia N, Lachiewicz A, et al. Advances in the treatment of fragile X syndrome. Pediatrics. 2009; 123(1): p. 378–90.

3. Darnell JC, Van Driesche SJ, Zhang C, Hung KY, Mele A, Fraser CE, et al. FMRP stalls ribosomal translocation on mRNAs linked to synaptic function and autism. Cell. 2011; 146(2): p. 247–61.

4. Ashley CT, Jr., Wilkinson KD, Reines D, and Warren ST. FMR1 protein: conserved RNP family domains and selective RNA binding. Science. 1993; 262(5133): p. 563–6.

5. Stoppel LJ, Kazdoba TM, Schaffler MD, Preza AR, Heynen A, Crawley JN, et al. RBaclofen Reverses Cognitive Deficits and Improves Social Interactions in Two Lines of 16p11.2 Deletion Mice. Neuropsychopharmacology. 2017.

6. Berry-Kravis EM, Lindemann L, Jonch AE, Apostol G, Bear MF, Carpenter RL, et al. Drug development for neurodevelopmental disorders: lessons learned from fragile X syndrome. Nat Rev Drug Discov. 2017.

7. Dolen G, Osterweil E, Rao BS, Smith GB, Auerbach BD, Chattarji S, et al. Correction of fragile X syndrome in mice. Neuron. 2007; 56(6): p. 955–62.

8. Qin M, Kang J, Burlin TV, Jiang C, and Smith CB. Postadolescent changes in regional cerebral protein synthesis: an in vivo study in the FMR1 null mouse. J Neurosci. 2005; 25(20): p. 5087–95.

9. Muddashetty RS, Kelic S, Gross C, Xu M, and Bassell GJ. Dysregulated metabotropic glutamate receptor-dependent translation of AMPA receptor and postsynaptic density-95 mRNAs at synapses in a mouse model of fragile X syndrome. J Neurosci. 2007; 27(20): p. 5338–48.

10. Bear MF, Huber KM, and Warren ST. The mGluR theory of fragile X mental retardation. Trends Neurosci. 2004; 27(7): p. 370–7.

11. Osterweil EK, Krueger DD, Reinhold K, and Bear MF. Hypersensitivity to mGluR5 and ERK1/2 leads to excessive protein synthesis in the hippocampus of a mouse model of fragile X syndrome. J Neurosci. 2010; 30(46): p. 15616–27.

12. Wang X, Snape M, Klann E, Stone JG, Singh A, Petersen RB, et al. Activation of the extracellular signal-regulated kinase pathway contributes to the behavioral deficit of fragile x-syndrome. J Neurochem. 2012; 121(4): p. 672–9.

13. Michalon A, Sidorov M, Ballard TM, Ozmen L, Spooren W, Wettstein JG, et al. Chronic Pharmacological mGlu5 Inhibition Corrects Fragile X in Adult Mice. Neuron. 2012; 74(1): p. 49–56.

14. Stoppel LJ, Osterweil EK, and Bear MF, The mGluR Theory From Mice to Men, in Fragile X Syndrome: From Genetics to Targeted Treatment Willemsen R and Kooy F, Editors. 2017 Elsevier.

15. Osterweil EK, Chuang SC, Chubykin AA, Sidorov M, Bianchi R, Wong RK, et al. Lovastatin corrects excess protein synthesis and prevents epileptogenesis in a mouse model of fragile X syndrome. Neuron. 2013; 77(2): p. 243–50.

16. Bradford RH, Downton M, Chremos AN, Langendorfer A, Stinnett S, Nash DT, et al. Efficacy and tolerability of lovastatin in 3390 women with moderate hypercholesterolemia. Ann Intern Med. 1993; 118(11): p. 850–5.

17. Mendola CE and Backer JM. Lovastatin blocks N-ras oncogene-induced neuronal differentiation. Cell Growth Differ. 1990; 1(10): p. 499–502.

18. Schafer WR, Kim R, Sterne R, Thorner J, Kim SH, and Rine J. Genetic and pharmacological suppression of oncogenic mutations in ras genes of yeast and humans. Science. 1989; 245(4916): p. 379–85.

19. Sebti SM, Tkalcevic GT, and Jani JP. Lovastatin, a cholesterol biosynthesis inhibitor, inhibits the growth of human H-ras oncogene transformed cells in nude mice. Cancer Commun. 1991; 3(5): p. 141–7.

20. Li W, Cui Y, Kushner SA, Brown RA, Jentsch JD, Frankland PW, et al. The HMGCoA reductase inhibitor lovastatin reverses the learning and attention deficits in a mouse model of neurofibromatosis type 1. Curr Biol. 2005; 15(21): p. 1961–7.

21. Asiminas A, Jackson AD, Louros SR, Till SM, Dando O, Bear MF, et al. Sustained correction of associative learning deficits following brief, early treatment in a rat model of Fragile X Syndrome. in preparation. 2018.

22. Caku A, Pellerin D, Bouvier P, Riou E, and Corbin F. Effect of lovastatin on behavior in children and adults with fragile X syndrome: an open-label study. Am J Med Genet A. 2014; 164a (11): p. 2834–42.

23. Pellerin D, Caku A, Fradet M, Bouvier P, Dube J, and Corbin F. Lovastatin corrects ERK pathway hyperactivation in fragile X syndrome: potential of platelet’s signaling cascades as new outcome measures in clinical trials. Biomarkers. 2016: p. 1–12.

24. Neuvonen PJ, Backman JT, and Niemi M. Pharmacokinetic comparison of the potential over-the-counter statins simvastatin, lovastatin, fluvastatin and pravastatin. Clin Pharmacokinet. 2008; 47(7): p. 463–74.

25. Tsuji A, Saheki A, Tamai I, and Terasaki T. Transport mechanism of 3-hydroxy-3-methylglutaryl coenzyme A reductase inhibitors at the blood-brain barrier. J Pharmacol Exp Ther. 1993; 267(3): p. 1085–90.

26. Krab LC, de Goede-Bolder A, Aarsen FK, Pluijm SM, Bouman MJ, van der Geest JN, et al. Effect of simvastatin on cognitive functioning in children with neurofibromatosis type 1: a randomized controlled trial. Jama. 2008; 300(3): p. 287–94.

27. Alabama-Birmingham U and NCI. A Randomized Placebo-Controlled Study of Lovastatin in Children With Neurofibromatosis Type 1 (STARS). ClinicalTrials.gov [Internet]. 2009-[cited 2010](ClinicalTrials.gov identifier: NCT00853580).

28. van der Vaart T, Plasschaert E, Rietman AB, Renard M, Oostenbrink R, Vogels A, et al. Simvastatin for cognitive deficits and behavioural problems in patients with neurofibromatosis type 1 (NF1-SIMCODA): a randomised, placebo-controlled trial. Lancet Neurol. 2013; 12(11): p. 1076–83.

29. Bearden CE, Hellemann GS, Rosser T, Montojo C, Jonas R, Enrique N, et al. A randomized placebo-controlled lovastatin trial for neurobehavioral function in neurofibromatosis I. Ann Clin Transl Neurol. 2016; 3(4): p. 266–79.

30. Payne JM, Barton B, Ullrich NJ, Cantor A, Hearps SJ, Cutter G, et al. Randomized placebo-controlled study of lovastatin in children with neurofibromatosis type 1. Neurology. 2016; 87(24): p. 2575–2584.

31. Thomson SR, Seo SS, Barnes SA, Louros SR, Muscas M, Dando O, et al. Cell-Type-Specific Translation Profiling Reveals a Novel Strategy for Treating Fragile X Syndrome. Neuron. 2017; 95(3): p. 550–563 e5.

32. Mans RA, Chowdhury N, Cao D, McMahon LL, and Li L. Simvastatin enhances hippocampal long-term potentiation in C57BL/6 mice. Neuroscience. 2010; 166(2): p. 435–44.

33. Lim JH, Lee JC, Lee YH, Choi IY, Oh YK, Kim HS, et al. Simvastatin prevents oxygen and glucose deprivation/reoxygenation-induced death of cortical neurons by reducing the production and toxicity of 4-hydroxy-2E-nonenal. J Neurochem. 2006; 97(1): p. 140–50.

34. Johnson-Anuna LN, Eckert GP, Franke C, Igbavboa U, Muller WE, and Wood WG. Simvastatin protects neurons from cytotoxicity by up-regulating Bcl-2 mRNA and protein. J Neurochem. 2007; 101(1): p. 77–86.

35. Proud CG. Signalling to translation: how signal transduction pathways control the protein synthetic machinery. Biochem J. 2007; 403(2): p. 217–34.

36. Kloog Y, Cox AD, and Sinensky M. Concepts in Ras-directed therapy. Expert Opin Investig Drugs. 1999; 8(12): p. 2121–2140.

37. Basso AD, Mirza A, Liu G, Long BJ, Bishop WR, and Kirschmeier P. The farnesyl transferase inhibitor (FTI) SCH66336 (lonafarnib) inhibits Rheb farnesylation and mTOR signaling. Role in FTI enhancement of taxane and tamoxifen anti-tumor activity. J Biol Chem. 2005; 280(35): p. 31101–8.

38. Clark GJ, Kinch MS, Rogers-Graham K, Sebti SM, Hamilton AD, and Der CJ. The Ras-related protein Rheb is farnesylated and antagonizes Ras signaling and transformation. J Biol Chem. 1997; 272(16): p. 10608–15.

39. Musumeci SA, Bosco P, Calabrese G, Bakker C, De Sarro GB, Elia M, et al. Audiogenic seizures susceptibility in transgenic mice with fragile X syndrome. Epilepsia. 2000; 41(1): p. 19–23.

40. Berry-Kravis E. Epilepsy in fragile X syndrome. Dev Med Child Neurol. 2002; 44(11): p. 724–8.

41. Yan QJ, Rammal M, Tranfaglia M, and Bauchwitz RP. Suppression of two major Fragile X Syndrome mouse model phenotypes by the mGluR5 antagonist MPEP. Neuropharmacology. 2005; 49(7): p. 1053–66.

42. Busquets-Garcia A, Gomis-Gonzalez M, Guegan T, Agustin-Pavon C, Pastor A, Mato S, et al. Targeting the endocannabinoid system in the treatment of fragile X syndrome. Nat Med. 2013; 19(5): p. 603–7.

43. King MK and Jope RS. Lithium treatment alleviates impaired cognition in a mouse model of fragile X syndrome. Genes Brain Behav. 2013; 12(7): p. 723–31.

44. Xie C, Sun J, Qiao W, Lu D, Wei L, Na M, et al. Administration of simvastatin after kainic acid-induced status epilepticus restrains chronic temporal lobe epilepsy. PLoS One. 2011; 6(9): p. e24966.

45. Yan QJ, Asafo-Adjei PK, Arnold HM, Brown RE, and Bauchwitz RP. A phenotypic and molecular characterization of the fmr1-tm1Cgr fragile X mouse. Genes Brain Behav. 2004; 3(6): p. 337–59.

46. Ling Q and Tejada-Simon MV. Statins and the brain: New perspective for old drugs. Prog Neuropsychopharmacol Biol Psychiatry. 2016; 66: p. 80–6.

47. Ramirez C, Tercero I, Pineda A, and Burgos JS. Simvastatin is the statin that most efficiently protects against kainate-induced excitotoxicity and memory impairment. J Alzheimers Dis. 2011; 24(1): p. 161–74.

48. Liao JK and Laufs U. Pleiotropic effects of statins. Annu Rev Pharmacol Toxicol. 2005; 45: p. 89–118.

49. Nurenberg G and Volmer DA. The analytical determination of isoprenoid intermediates from the mevalonate pathway. Anal Bioanal Chem. 2012; 402(2): p. 671–85.

50. Istvan E. Statin inhibition of HMG-CoA reductase: a 3-dimensional view. Atherosclerosis Supplements. 2003; 4(1): p. 3–8.

